# Formation Of Small-World Network Containing Module Networks In Globally And Locally Coupled Map System With Changes In Global Connection With Time Delay Effects

**DOI:** 10.1101/2022.11.13.516347

**Authors:** Taito Nakanishi, Akinori Awazu

## Abstract

In this study, we performed comprehensive morphological investigations of spontaneously formed network structures among elements in coupled map systems involving global connections that change depending on the synchronicity of states of elements and spatially local connections. The model formed various hierarchical networks, some of which were classified as small-world networks containing multiple module networks, similar to the neural network of mammalian brains. Moreover, such complex networks were formed in wider parameter regions when the global connection to an element from the other element was strengthened by the synchronization between the present and past states of the former and latter elements, respectively. This study suggests that the time delay effects for connection changed among elements and local interactions promoted the self-organization of small-world networks containing module networks, such as neural networks; neural networks contain them as spike-timing-dependent plasticity and inter-neuron interaction through glial cells.

## 1. Introduction

Various biological and social systems are regulated via their self-organized network structures. Neural networks [1-17], metabolic networks [18-21], food webs [22,23], and human communities [24-29] are typical networks that exhibit self-regulation of connectivity through learning, cell differentiation and adaptation, evolution, and communication.

The self-regulation behaviors of such systems have been studied using models that comprise dynamic elements involving changes in mutual relationships among the elements. For example, models of neural networks comprising elements that imitate a nerve cell have been described as excitable or chaotic oscillators. Consequently, the temporal change in coupling among these elements was assumed to follow a rule inspired by Hebb’s law or spike-timing-dependent plasticity (STDP) [1–17]. Recent studies have suggested that one of the typical classes of such models, coupled chaotic map systems with Hebbian-like rules, may frequently exhibit spontaneous formations of asymmetrically connected network structures with the emergence of one-way hierarchies and/or loops, multiple layers, and pacemakers [5-7,16,17].

Recent social and medical studies have suggested that such self-organized networks often form small-world networks with specific statistical properties in the edge distributions among nodes. For example, small-world networks have been found in various human social networks [30]. Neural networks in mammalian brains are known to form small-world networks [31,32], which also involve multiple module networks [33-35]. Recent mathematical studies have suggested simple procedures for painting small-world networks [30]. However, recently proposed dynamic network system models involving the self-regulation of their connection topologies did not provide sufficient mechanisms for the manner in which such social and neural networks could organize small-world networks simultaneously and robustly.

Most recent mathematical models for the studies of self-organized networks were constructed based on globally coupled dynamical systems with temporally changeable interaction strengths among elements. However, social networks appear to involve global, distance-free interactions such as those through world-wide web systems and local interactions such as communication in family, among friends and coworkers, and in various local communities. In addition, neural networks in the mammalian brain involve both the global synaptic interactions among neurons and local interactions among spatially neighboring neurons through glial cells occupying spaces among neurons; the volume fractions of glial cells are 2–10 times more than those of neurons [36-41]. Thus, the combination of local and global interactions with temporally changeable interaction strength may be considered to promote the self-organization of small-world networks.

In this study, we developed dynamic systems comprising spatially distributed elements connected globally and locally. All elements were connected globally with temporal changes in connection strengths among all elements, and neighboring elements were connected locally with constant strength. Subsequently, through comprehensive simulations, the abovementioned conjectures were examined for the self-organization and robust formation of small-world networks with multiple module networks.

## 2. Model and Methods

### 2.1 Model

As a simple dynamical system with local and global interactions exhibiting element state-dependent changes in global interaction strength, the following coupled map system with N elements was considered:

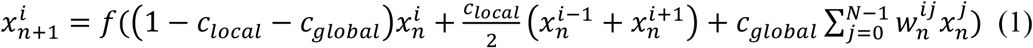

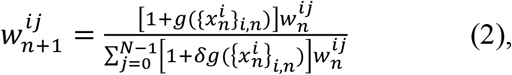

where 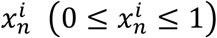 and 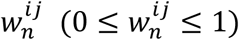 are the state of the *i*-th element (*i* = 0, 1, … *N* − 1) and the connection strength from the *j*-th to *i*-th element at time *n*, respectively, and *f*(*x*) = *ax*(1 − *x*) (logistic map) is assumed. We assumed that the elements were spatially arranged in a circle in the order *i* = 0, 1, …, n − 1. Further, it was assumed that the 0-th element and the (n − 1)-th element were adjacent to each other and provided *i* + 1 = 0 for *i* = (*N* − 1)-th element and for *i* = 0-th element. The first, second, and third terms in function *f* in Eq. (1) indicates the influence of each element itself, that of two spatially neighboring elements, and that of globally coupled elements. In this model, *w*^*ii*^ = 0 (no self-connection) was assumed, and the condition of 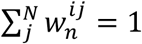 was always satisfied for *i*. The parameters *a, c*_*local*_, and *c*_*global*_ indicate the parameters of the logistic map, strength of local interaction between two spatially neighboring elements, and strength of the influences of global interaction, respectively.

In the case of 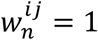, the present model is equal to the coupled map model with local couplings and uniform global couplings proposed by Ouchi and Kaneko, showing various complex spatio-temporal pattern dynamics [42]. When 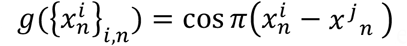 with *c*_*local*_ = 0 and that of 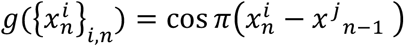 with *c*_*local*_ = 0, the present model is equal to that proposed by Ito and Ohira [5]. Here, in the former case, the connection from *j* -th to *i* -th element changes depending on the synchronicity between the states of *i*-th and *j*-th elements. Whereas, in the latter case, the changes depend on the synchronicity between the present state of *i*-th element and the one-step past state of *j*-th element. These models showed a rich variety of self-organizations of directionally connected networks with multiple layers, hierarchies, and/or loops, and a pacemaker community in an exhaustive study of these models [16, 17].

In the present model, the set 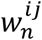 represents the directional network structures of the elements at time *n*. Herein, the connection from the *j*-th to the *i*-th element is considered strong (weak) when 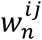 is large (small). In general, 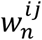 changes in *n*. Therefore, as employed in previous studies [4-7,9,16,17], we first classified the connection profiles of the typical networks obtained via the proposed model according to the following definition: the connection from *j*-th to *i*-th elements exists if the time average of 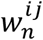, ⟨*w*^*ij*^⟩, satisfied 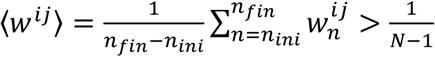, where 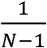 indicates the average of 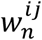 over the entire set of *i* and *j* (*i* ≠ *j*). This study referred to the network characterized by the set of ⟨*w*^*ij*^⟩ as the effective network.

### 2.2 Simulation method

We performed simulations of the model for each set of *a, c*_*local*_, and *c*_*global*_ for five different initial conditions, and focused on the most frequently obtained network structures with *n*_*ini*_ = 10^6^ and *n*_*fin*_ = 2 × 10^6^ as the typical network structure. As the initial condition of each simulation, 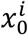 was chosen randomly from the values satisfying 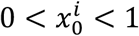, and 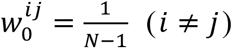 was provided, assuming that all elements were uniformly connected to all the elements, except for itself. We confirmed that the influence of the initial conditions was negligible when *n*_*ini*_ = 10^6^ was chosen. This is supported by the relaxation behaviors of the autocorrelation functions of the connection strength among the elements described as

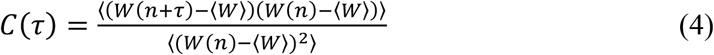

where 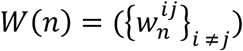 is an *N*(*N* − 1) dimensional vector. Here, < *W* > was assumed to be 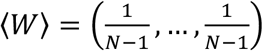 because the average of 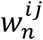 was estimated to be 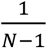 if 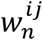 changed ergodically in *n*.

### 2.3 Estimations of small world propensity and modularity of “undirected networks”

Neural networks in mammalian brains are typical network systems with element state-dependent connection changes. Undirected networks described from neural networks in mammalian brains can form small-world networks [31,32]. Additionally, these neural networks contain multiple module networks where neurons are divided into modules, and intra-module connections among neurons are dense, whereas inter-module connections are sparse [33-35].

Thus, to evaluate the structural features of the formed effective networks, we measured the small-world propensity (SWP), modularity, and number of modules of each network to compare the statistical similarities between the networks obtained from the present model and the neural networks in the mammalian brain as follows. Modularity is defined as a value that increases with the differences between the connection number among elements in the same modules and those among elements in different modules [33].

To evaluate these statistical values for each network, we first defined the undirected connectivity between *i* -th and *j* -th elements as 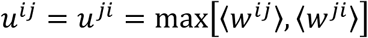, and connections satisfying 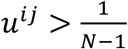 as “undirected connections”. In addition, we defined the “undirected effective network” as that comprising connections with 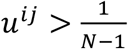 for each network.

Second, the SWP, modularity, and number of modules of each undirected effective network obtained by each parameter set were evaluated based on the following recently proposed methods. The SWP was calculated according to the method proposed by Watts and Strogatz [30,31], which was employed to evaluate the neural network topology of mammalian brains [31,32]. Whereas, the modularity and number of modules in each network were estimated using the method proposed by Newman et al [33,43], which was also employed to extract the module structures of the neural networks [33-35]. The network was regarded as a small-world network when the SWP > 0.6, and it was expected to contain certain modules when modularity > 0. In recent studies on neural networks in mammalian brains, the SWP was estimated 0.65 ∼ 0.9 [31,32], and positive modularity was obtained [33-35].

## 3. Results

### 3.1 Influence of local connections and time delay effects on global connection changes in formed effective network structures

To reveal the influence of the local connections and the time delay effects on connection changes in the formed network structures, we simulated the model with I) 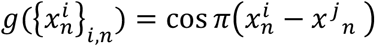 and *c*_*local*_= 0, II) 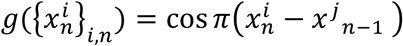 and *c*_*local*_ = 0, III) 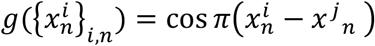 and *c*_*local*_ > 0, and IV) 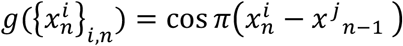 and *c*_*local*_ > 0. These models were referred to as I) model SS: considered only the effects of simple synchronization among elements, II) model SD: considered the effects of synchronization and time delay among elements, III) model SSL: considered the effects of simple synchronization and local connections among elements, and IV) model SDL: considered the effects of synchronization, time delay, and local connections among elements.

Figure 1 shows the phase diagrams of typical effective networks obtained using the abovementioned four models with *N* = 30 as a function of *a* and *c*_*global*_. Here, the case of *c*_*local*_ = 0.1 is shown for model SSL, and the cases of *c*_*local*_ = 0.1, *c*_*local*_ = 0.05, and *c*_*local*_ = 0.2 are shown for model SDL. Each symbol in Fig. 1 represents the mean of the typical network structures observed for a given parameter set. The phase diagrams of the SS models shown in Fig. 1(a) exhibited the same results as those reported in a previous study [17]. Additionally, although the symbols used to indicate each formed network were different, the phase diagrams of the SD models shown in Fig. 1(c) exhibited the same results as those reported in another previous study [16] for *c*_*global*_ ≥ 0.1. The results of *c*_*global*_ = 0.05 are shown in Fig. 1(c). Furthermore, the behaviors of the SSL and SDL models for *c*_*global*_ > 0.3 were obtained as similar to those observed in the parameter regions marked U, G, and P, as described later.

**Fig. 1.**
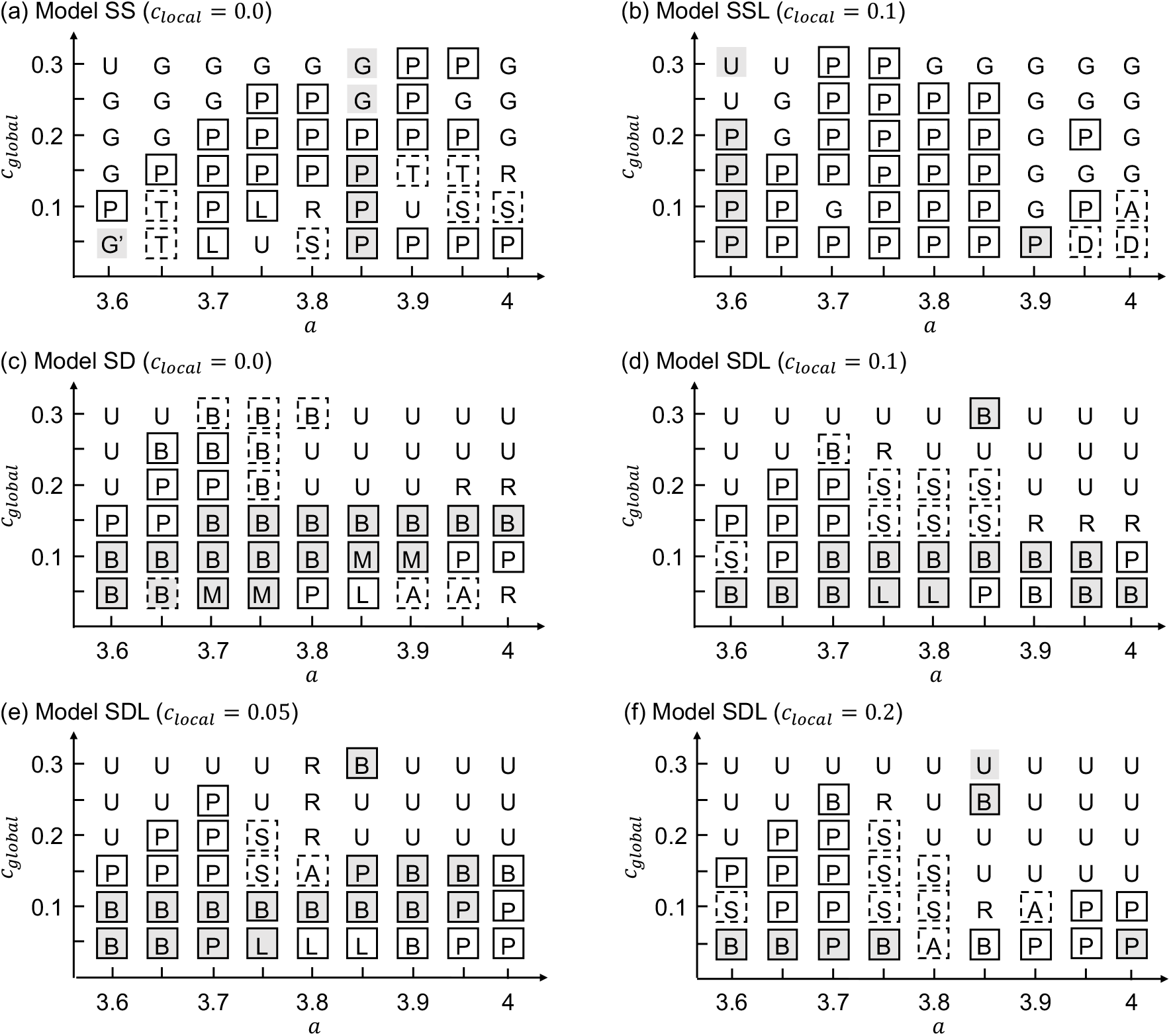
Phase diagrams of the typical effective networks obtained by the models (a) SS, (b) SSL with *c*_*local*_ = 0.1, (c) SD, and (d-f) SDL with *c*_*local*_ = 0.1, *c*_*local*_ = 0.05, and *c*_*local*_ = 0.2. Each symbol indicates each formed effective network (details in test). Rigid box indicates the connections among elements 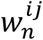 are stationary, and broken box indicates that connections 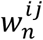 changed temporally. Shadowed background indicates that the states of all elements 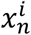 exhibit periodic changes in time, whereas white background indicates that the states of certain or all elements exhibit chaotic motions.

The temporal change in 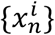 exhibited periodic or chaotic motion depending on the parameter set, as shown in Fig. 1. In the parameter regions marked U and G, the following simple trivial networks were obtained, where the elements were effectively connected symmetrically and uniformly (partially uniformly) with each other, and the parameter regions marked U, 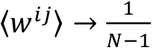 and 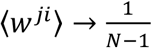 were obtained. In the parameter regions marked as G, the elements were divided into groups such that 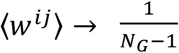 and 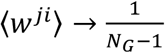 were obtained in each group, where *N*_*G*_ indicates the number of elements in each group. In the subsequent arguments, we do not focus on simple, trivial cases.

For other parameter regions, various networks with different structural characteristics were obtained as follows. The typical example structures of the effective networks in the case of *N* = 30 are shown in Fig. 2.

**Fig. 2.**
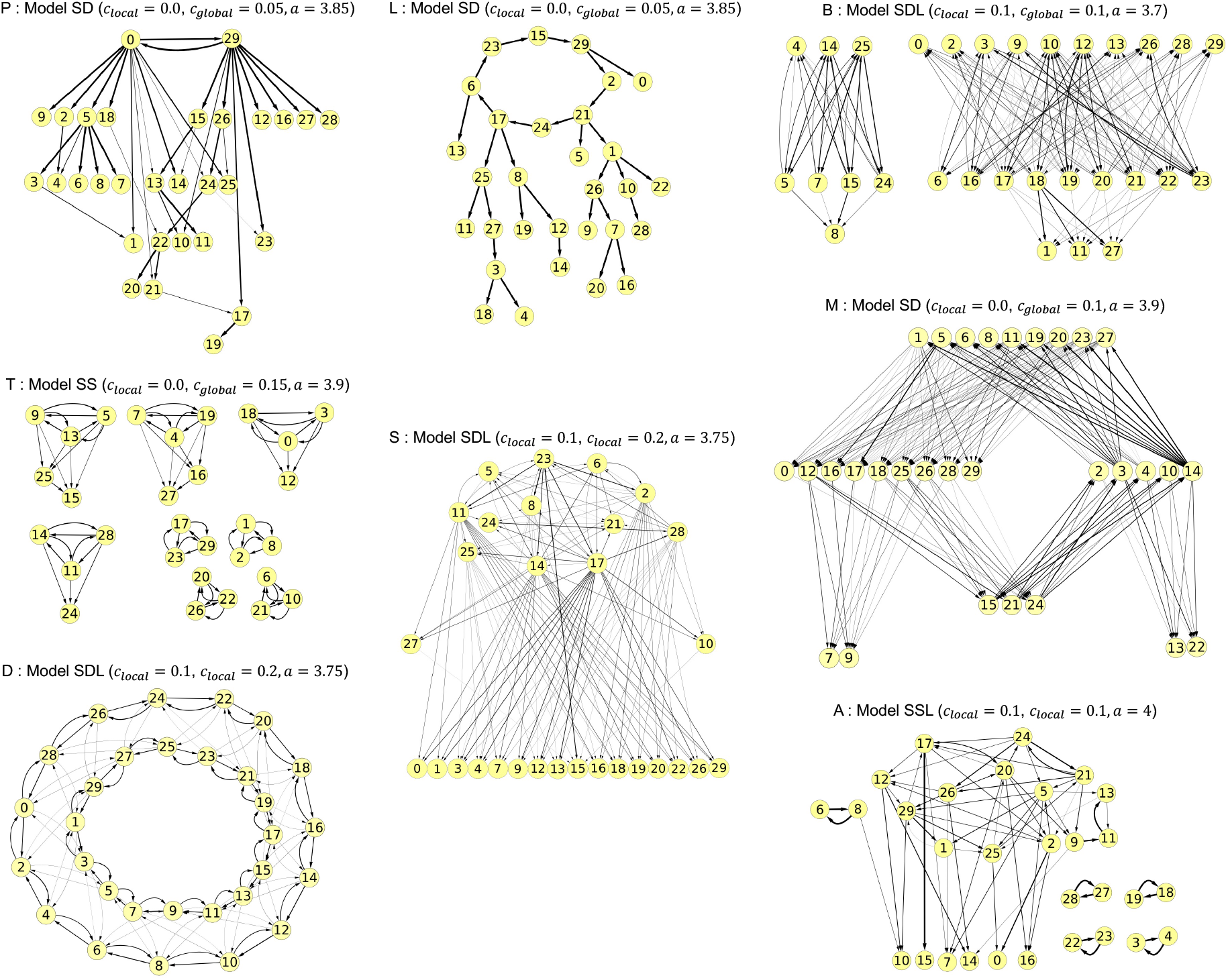
Typical structures of effective networks for respective symbols in Fig. 1. Each circle with index *i* indicates *i*-th element, and the arrow from the *i*-th circle to the *j*-th one indicates existence of a connection from *i*-th element to *j*-th one. Index *i* also indicates its spatial position (details in text).

P: Pair-driven network, where certain elements form pairs that are connected symmetrically with each other, whereas the other elements form one-way directionally connected networks without any loops.

L: Loop-driven network, where a few elements are located in the upper stream of the network and form a one-way connected loop, whereas the other elements form one-way directionally connected networks without any loops.

T: Trio-driven networks, where certain elements form trios, which are connected almost symmetrically to each other, whereas the other elements form one-way directionally connected networks without any loops.

S: Sym-community-driven networks where some elements form a community, whereas the other elements form one-way directionally connected networks without any loops. The elements in the community are connected almost symmetrically, and each element in this community is connected to certain non-community-forming elements. These networks are large hierarchical networks containing one such community in their uppermost stream.

B: Bipartite layer-driven networks, where certain elements form paired layers upstream of the network. Each element in one upstream layer is connected to and from other elements that belong to another upstream layer. Further, certain elements in these layers are connected to non-layer-forming elements.

M: Multilayer networks, where the elements are divided into more than three layers, and each element is connected to and from other elements that belong to other layers. Here, certain layers are located in the upper stream of the network and form a one-way connected loop, whereas the other elements form one-way directionally connected networks without any loops at the lower stream of the network.

The detailed features of the abovementioned networks, P, L, T, and S, have already been reported in a previous study [17], and those of networks B and M have also been reported [16]. Conversely, the following network structures have not been reported in previous studies:

D: Double-strand network, where 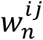 changes chaotically in *n*, and ⟨*w*^*ij*^⟩ with 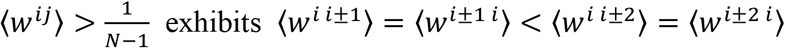.

A: Asym-community-driven network, where certain elements formed a community upstream of the network, similar to a sym-community-driven network. However, the elements in the community were connected almost asymmetrically to each other, unlike a sym-community-driven network.

Comparisons among the phase diagrams of the four models showed that the local connections appeared to expand the parameter region to form a pair-driven network (network P) and a sym-community-driven network (network S). In particular, the SSL model exhibited network P in most cases, whereas both P and S networks emerged in the SDL model. In addition, the parameter region wherein networks P or S emerged was wider with an increase in *c*_*local*_. Moreover, the SS model could also exhibit network S; however, the SDL model showed network S in a wider parameter region than SS model.

### 3.2 Characterizations of formed effective networks by small world propensity and modularity of undirected effective networks

To evaluate the structural features of effective networks, the SWP and modularity were estimated for undirected effective networks obtained using each parameter in each model. Figure 3 shows (a) SWP as a function of *a* and *c*_*global*_, and (b) modularity as a function of *a* and *c*_*global*_, where *c*_*local*_ = 0.1 for models SSL and SDL. SWP was measured when networks P, L, T, S, B, M, A, or D were formed. Additionally, modularity was estimated only for the parameter region where the undirected effective network was regarded as a small-world network with an SWP larger than 0.6. In these figures, the parts of networks P and S, as shown in Fig. 4, exhibited a value of SWP larger than 0.6, and a value of modularity larger than 0. Additionally, the SDL model exhibited such effective networks in a wider range of parameter regions than that in the other models.

**Fig. 3.**
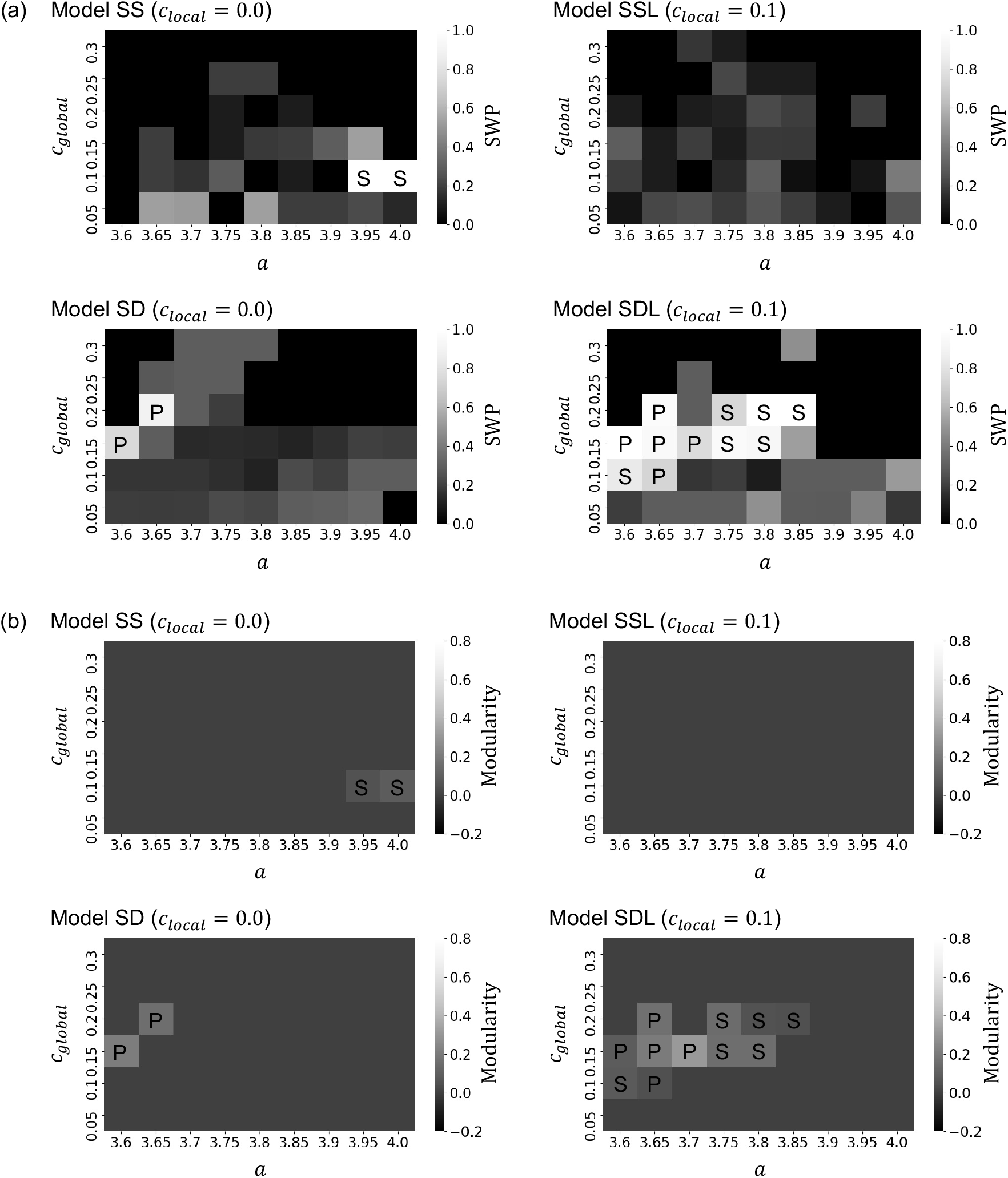
(a) Small world propensity (SWP) and (b) modularity measured by the models SS, SD, SSL with *c*_*local*_ = 0.1, and SDL with *c*_*local*_ = 0.1. SWP was put as 0 in (a) when the trivial network (U or G) was formed. Modularity is calculated only when SWP > 0.6 and is set as 0 when SWP ≤ 0.6 in (b).

**Fig. 4.**
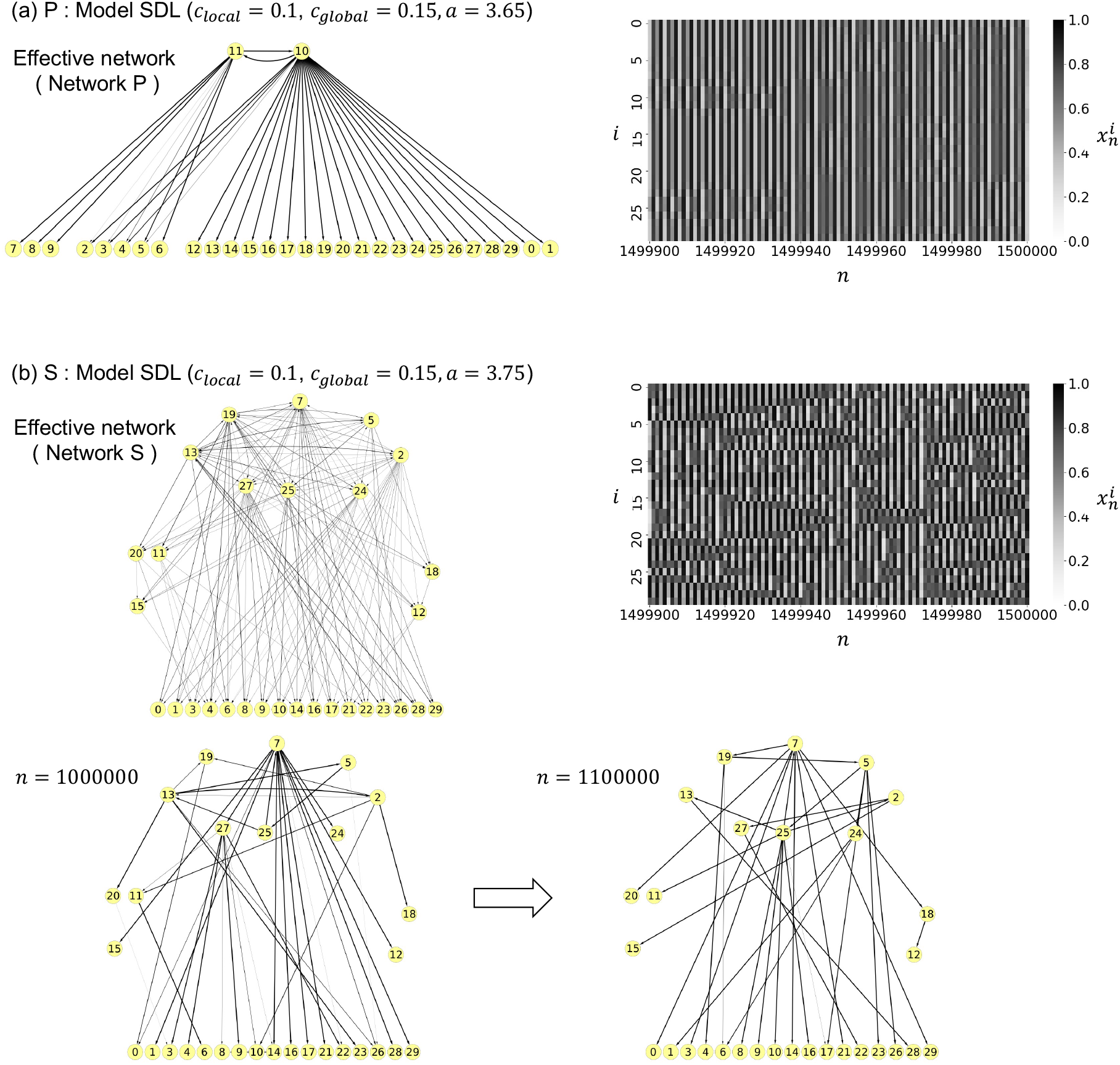
Typical structural features of (a) pair-driven networks P formed by model SDL with *a* = 3.65, *c*_*global*_ = 0.15, and *c*_*local*_ = 0.1, and (b) hidden community-driven networks S formed by model SDL with *a* = 3.75, *c*_*global*_ = 0.15, and *c*_*local*_ = 0.1, exhibiting small world propensity > 0.6 and positive modularity in cases of *N* = 30. The structure of network P is obtained as stationary for sufficiently large *n*, and then, the network at time *n* converges to effective network in (a). Conversely, the structure of network S changes in *n* even for sufficiently large *n*, where the obtained structures are different among the effective network (upper), the network at *n* = 1000000 (lower left), and that at *n* = 1100000 (lower right), as shown in (b).

### 3.3 Sym-community-driven networks exhibited similar small world propensity and modularity of undirected networks to neural network of mammalian brain

Certain pair-driven networks (network P) and sym-community-driven networks (network S) tend to exhibit high values of SWP and modularity. It was noted that both networks commonly involved a hierarchical structure, with an upper community of elements connected symmetrically and lower elements with few connections from them. However, in contrast to network P, network S contained intermediate elements containing connections from certain upper elements and those to certain lower elements. In addition, although the upper community in network P always comprised only two elements, the number of elements constructing the upper community in network S was generally greater than 2.

To evaluate the robustness of the abovementioned features, the system-size-dependent behaviors of the model SDL with the parameters of networks P or C were obtained. Figure 5 shows the averages and standard deviations of (a) SWP, (b) modularity, and (c) number of modules as a function of the number of elements *N* in the cases where networks P and S were formed; typical shapes of (d) network P and (e) network S were formed in the case of *N* = 120. The averages and standard deviations were calculated using simulations from 10 different initial conditions for each *N*. This figure showed the SWP was almost independent of *N*. However, the modularity and number of modules increased with *N* when the network S was formed. Whereas, the modularity decreased with *N* when the network P was formed. These facts suggest that the SDL model with appropriate parameters can form a larger small-world network with a multitude of modules according to the increase in the number of elements.

**Fig. 5.**
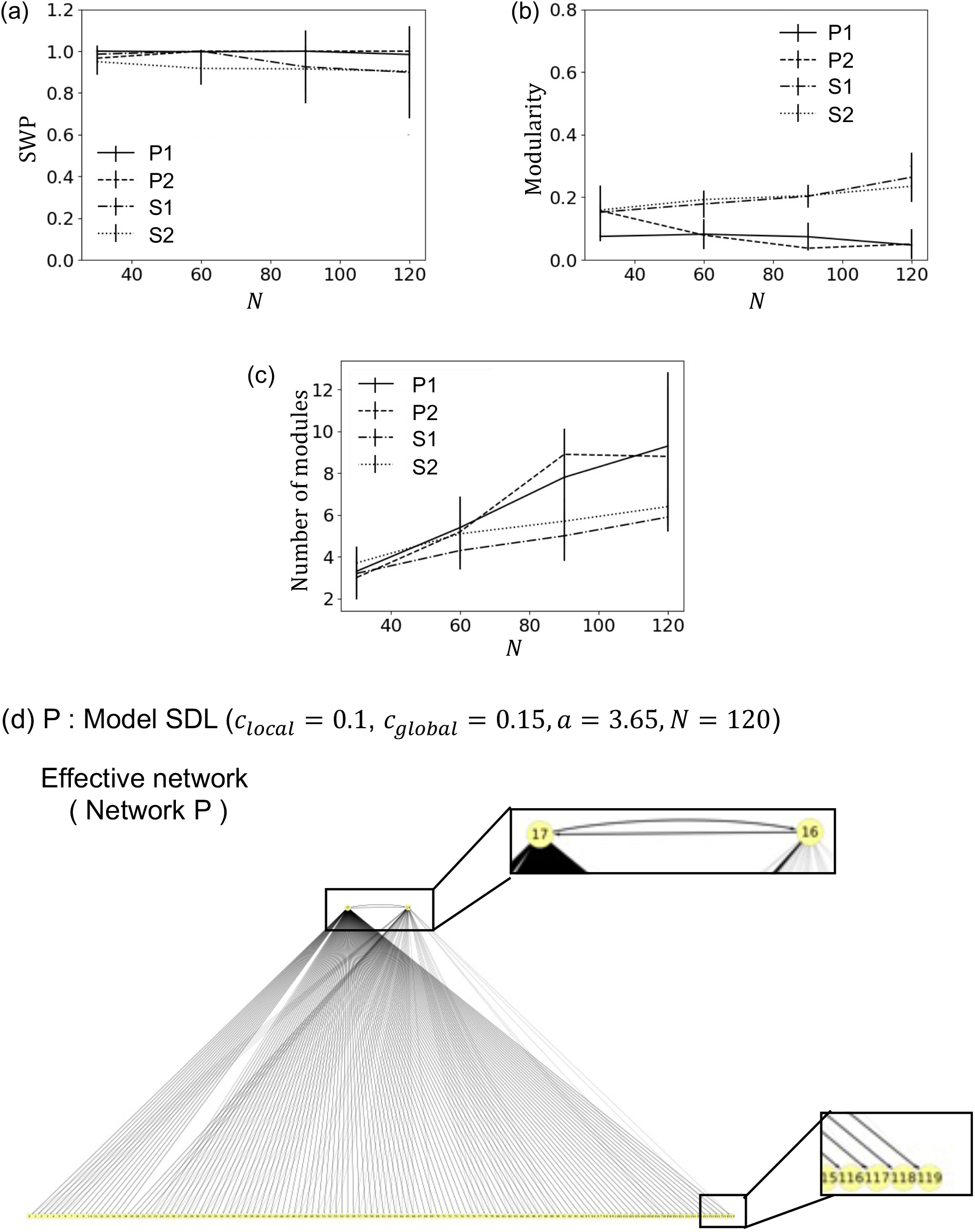
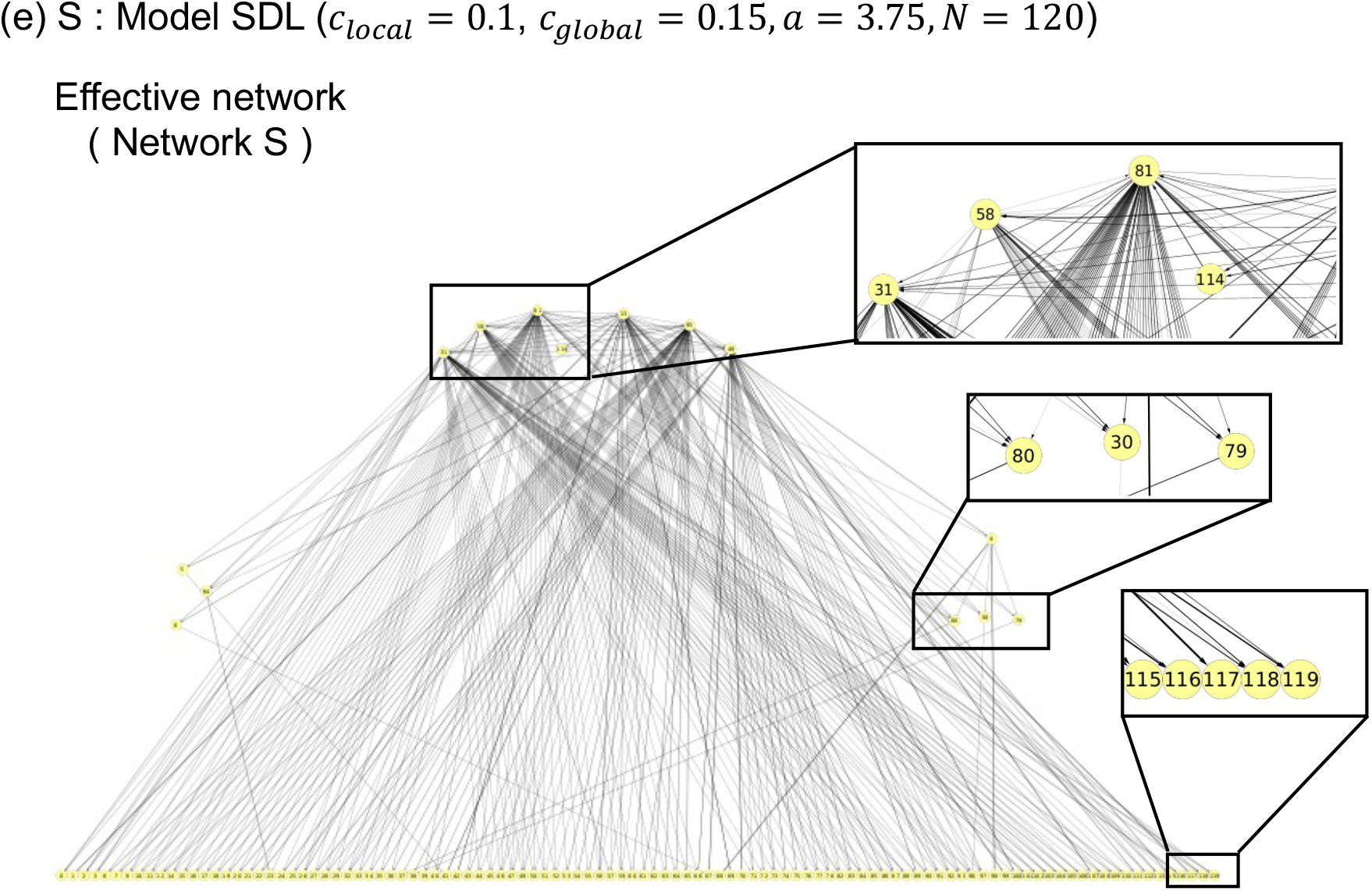
Average (curves) and standard deviation (error bar) of (a) small world propensity, (b) modularity, and (c) number of modules as a function of *N* when the network P or S are obtained by the SDL model. P1 and P2 indicate that the model exhibits the network P, where parameters are *a* = 0.36 (P1) and *a* = 0.365 (P2) with *c*_*global*_ = 0.1 and *c*_*local*_ = 0.15. S1 and S2 indicate that the model exhibits the network P, where parameters are *a* = 0.375 (S1) and *a* = 0.38 (S2) with *c*_*global*_ = 0.1 and *c*_*local*_ = 0.15. (d) Typical structures of networks P and (e) that of network S obtained as effective networks in the case of *N* = 120, where parameters are *a* = 0.365 in (d) and *a* = 0.375 in (e) with *c*_*global*_ = 0.1 and *c*_*local*_ = 0.15.

## 4. Discussion

In this study, we comprehensively focused on the morphologies of spontaneously formed network structures using globally and spatially (local) coupled map systems involving global connection changes dependent on the synchronicity of states of elements. Various types of hierarchical directional networks were obtained through the simulations of the proposed model, and some of them were classified as small-world networks containing multiple module networks, as observed in the neural network of mammalian brains. In addition, we found such small world networks were formed in wide range of parameter regions when the temporal change in each global connection strength between two elements was influenced by their present and past states.

The comparison with the results of the present arguments and the behaviors of neural networks in mammalian brains indicated the following common features:

I) The synaptic connections among neurons in the brain exhibited spike-timing-dependent-plasticity [44], indicating that the change in each global connection from one element to another in both systems was strengthened by the synchronization of the past and present states of the former and latter elements, respectively. II) The spatially neighboring neurons mutually influenced each other through the glial cells that occupied spaces among neurons [36-41], indicating that both systems involved interactions among spatially neighboring elements. III) Both systems can form a small-world network containing modular networks [31-35].

Conversely, the basic dynamics of each element were significantly different between the proposed model and neural networks in the brain when each element was isolated, where the element in the proposed model exhibited chaotic dynamics, whereas that in neural networks showed excitable dynamics. Thus, in the formation of small-world networks containing multiple modular networks, the effects of the time delay involved in the change in the connection strength to an element from other elements and the local interactions between spatially neighboring elements appeared to contribute more significantly than the detailed features of each element.

In this study, the question “Why and the manner in which the local interactions and the effects of the time-delay involved in the change in the global connection strengths promote the small world networks?” was not answered. To answer this question, additional theoretical studies should be conducted from the perspective of dynamical systems with large degrees of freedom in the future.

The present results are expected to provide novel insights into the self-organization of small-world networks by not only neural networks but also various other biological and social networks. To reveal the detailed processes and the mechanism of the formation of such biological networks, the construction of models or modifications of recently proposed models comprising more realistic model elements with interaction rules based on the present arguments should be conducted, which is also one of the future issues.

## Acknowledgments

We thank Junji Ito, Takahiro Chihara, Amika Ohara, and Masashi Fujii for providing fruitful information. This work was supported by Grants-in-Aid for Scientific Research (KAKENHI) [Grant Number 21K06124 (A.A.)] from the Japan Society for Promotion of Science. Computations were partially performed on the NIG supercomputer at ROIS National Institute of Genetics.

